# A simple computer vision algorithm as a clinical aid for the pathologist

**DOI:** 10.1101/853325

**Authors:** J.M. Lázaro-Guevara, B.J. Flores-Robles, A.E. Murga, K.M. Garrido

## Abstract

Histological analysis for cancer detection or stratification is performed by observing and examining a small portion of a biopsied tissue under a microscope. Nevertheless, to assign clinical meaning to the findings, the analysis and interpretation of an experienced Pathologist is always necessary. Using high-resolution images, these experts visually examine the sample looking for specific characteristics on the cell shapes and tissue distributions, so they could decide whether tissue regions are cancerous, and establish the malignancy level of it. However, with the increasing demand for work for those pathologists and the importance of accuracy on diagnostics, multiple attempts to simplify their work have been performed. Current Imaging technologies allow novel horizons in the automatized selection of some of the characteristics that indicate malignancy in a biopsy. In this work, we propose a simple computer vision algorithm that can be implemented as a screening method for focusing in histological areas with higher risk of malignancy saving time to the pathologist and helping to perform a more standardized work, an easy observation with the potential to become in an aid to daily clinical work.

## I. Introduction

Histology is the study of the microscopic anatomy of cells and tissues of organisms.

Histological analysis is performed by observing and examining a small portion of the biopsied tissue under an optical or electron microscope. But for a clinical meaning of this analysis the interpretation of an expert and experienced Pathologist is always necessary, in histology image analysis for cancer diagnosis, this expert visually examine some characteristics on the cell shapes and tissue distributions, so they could decide whether tissue regions are cancerous, and determine the malignancy level.

Such histopathological study has been extensively employed for cancer detection and grading applications, including prostate, breast, cervix, and lung, however the new technologies allow new horizons in the automatized selection of this characteristics for helping this experts to perform a more standardized work, an easy observation(1, 2).

This new horizon is aimed to find a way to detect cancer cells in histology segments by computer imaging; this has been the “Holy Grail” to surgeons for a quick and accurate response in the operating room. Now we have the Tele-Medicine, also the Tele-Pathology, but because the lack of trained pathologist in the moment of surgery, a computer vision program that can assess a good result discriminating between cancer and normal cell will be very helpful for determinate the borders amplitude in excision surgery.

Many attempts had be done for this objective as shown by (Lei et al, 2012), where described a way to achieve an automated carcinoma detection and classification(1)(3), even has been used Deep Neural Networks (4, 5), and histogram decomposition (6, 7), but still there is no software on market or in clinical usage that can achieve this goal better or at least equal than a trained pathologist (8, 9). In this work we attempted to create a tool that can be used by the pathologist as a screening method for focusing in histological areas with higher risk of malignancy saving time to the pathologist and becoming in a aid to daily clinical work.

## II. Methods

### A. Determination of the parameters to evaluate and type of Cancer

The five (5) most common parameters present in neoplasic tissues:

1. Citoplasmic – nucleus relationship
2. Alignment or cell orientation pattern
3. Increment of mitosis,
4. Increment on vasculature
5. Presence of necrosis

This study was focused on poorly differentiated adenocarcinoma stage 3 and 4, for the early stages of analysis in software. And the 5 parameters where selected as suggestion of the Department of Pathology of Hospital San Juan de Dios in Guatemala, after that suggestion the characteristics of each one were correlated with the information included in Robbins Basic Pathology, 9th Edition.

Also images for doing the test as a primary source, were proportioned by the Guatemala Pathology department, and as second source http://cancer.digitalslidearchive.net/, site that is free on access and allow to do a control over the type of images, to compare against as the primary source.

### B. PIXEL filtering and selection

First step was determinate the Citoplasmic nucleus relationship, for that purpose was created and histogram dispersion of each channels of an JPG image (RGB), in 50 histological images using hematoxylin-eosin to achieve and average of the ranges of each channel for nucleus (Purple) and cytoplasm (Pink) (figure 1). Obtaining the values for purple (R 70 – 150; G 60 – 100; B 145 – 255) and Pink (R 225 – 255; G 0 – 40; B 150 – 200).

**Figure 1.**
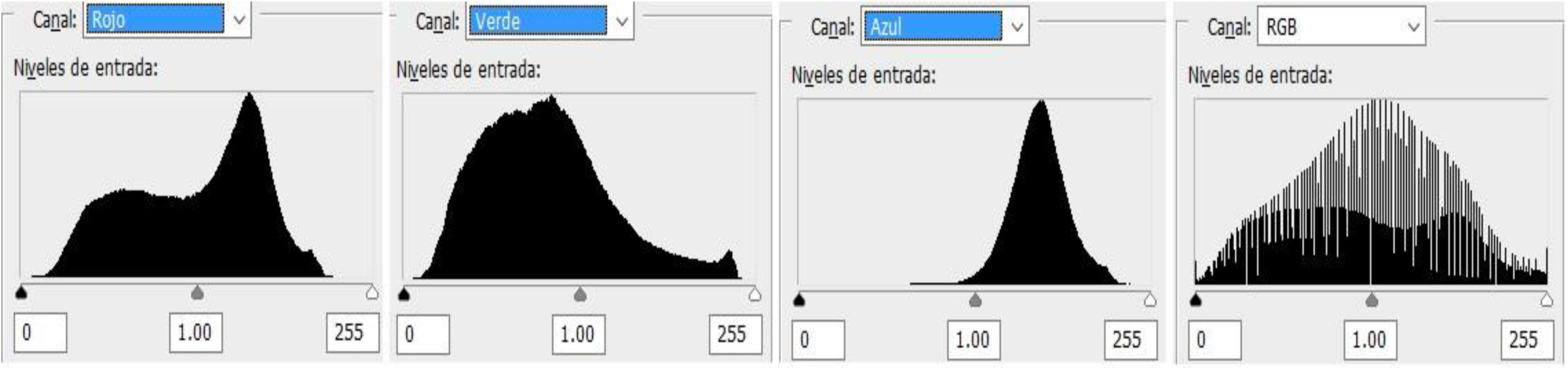

### C. Thresholding and edge detection for nuclei and cytoplasm (relationship)

Based on pixel information selection realized on step B, a trinary image was created using the zonal gray scale technique (figure 2a), (Purple =128; Pink=0; other=255), after that a binary thresholding algorithm, similar to Otsu’s method, and then subjected the output to Sobel Edge Detection where applicable, to obtain the inner and external borders of the cell and surroundings (figure 2b), it was previously applied a blurred technique and compare both images, the blurred one, and the normal, image to edge detection, obtained variable results, from improve in 20% or images to no difference, for that reason the blurring step is declared as optional, and not included in the images shown in the research.

**Figure 2.**
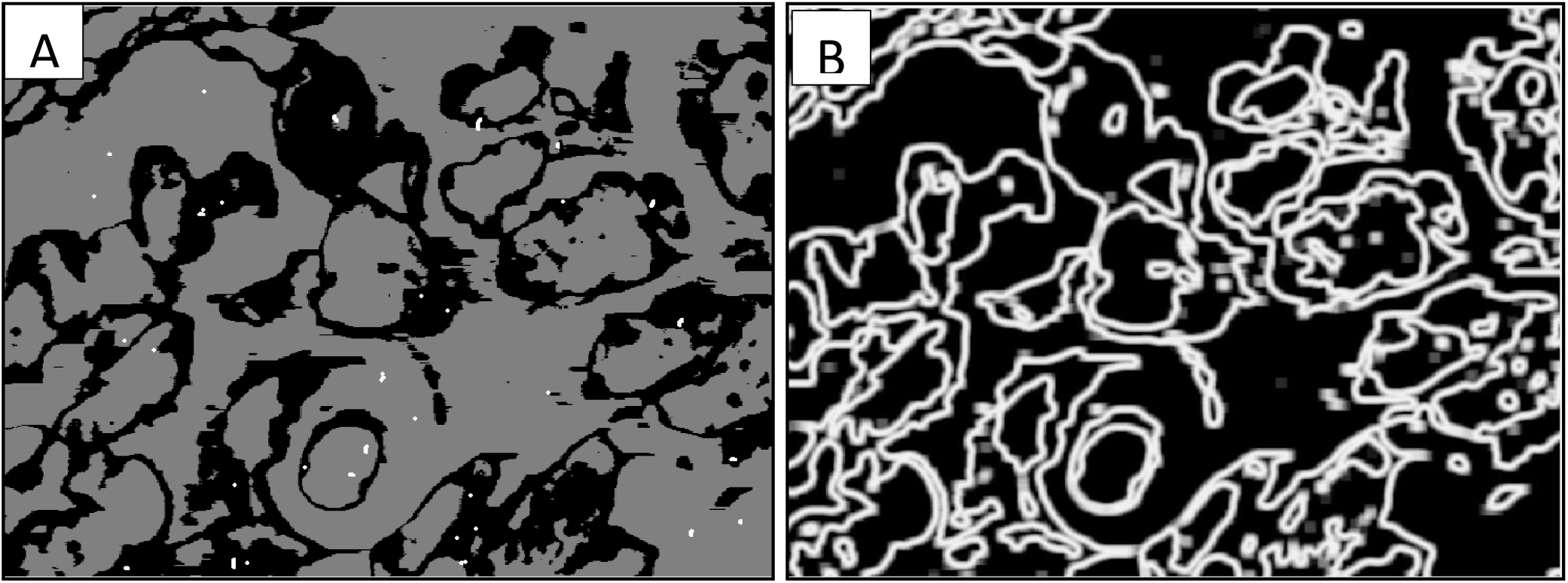

After detecting the borders, they were better delimited and defined as nucleus and cytoplasm, and high contrast applied filtering the Green channel on the image, also an index of Citoplams-Nucleus were created by two methods, area subtracting and pixel counting to fulfill the criteria, that a cancer cell the nucleus is major than the surrounding cytoplasm.

### D. Necrosis and Increment on vascularity detection

Using the same technique applied on step A for pixels filtering and knocking down the Blue channel with the remaining two channels filtered to (R >= 190) and (G <= 120), it is possible to obtain the vasa and vascularity information, for necrosis it is necessary to filter (R >= 210) and (G >= 200) as shown (figure 4).

**Figure 3.**
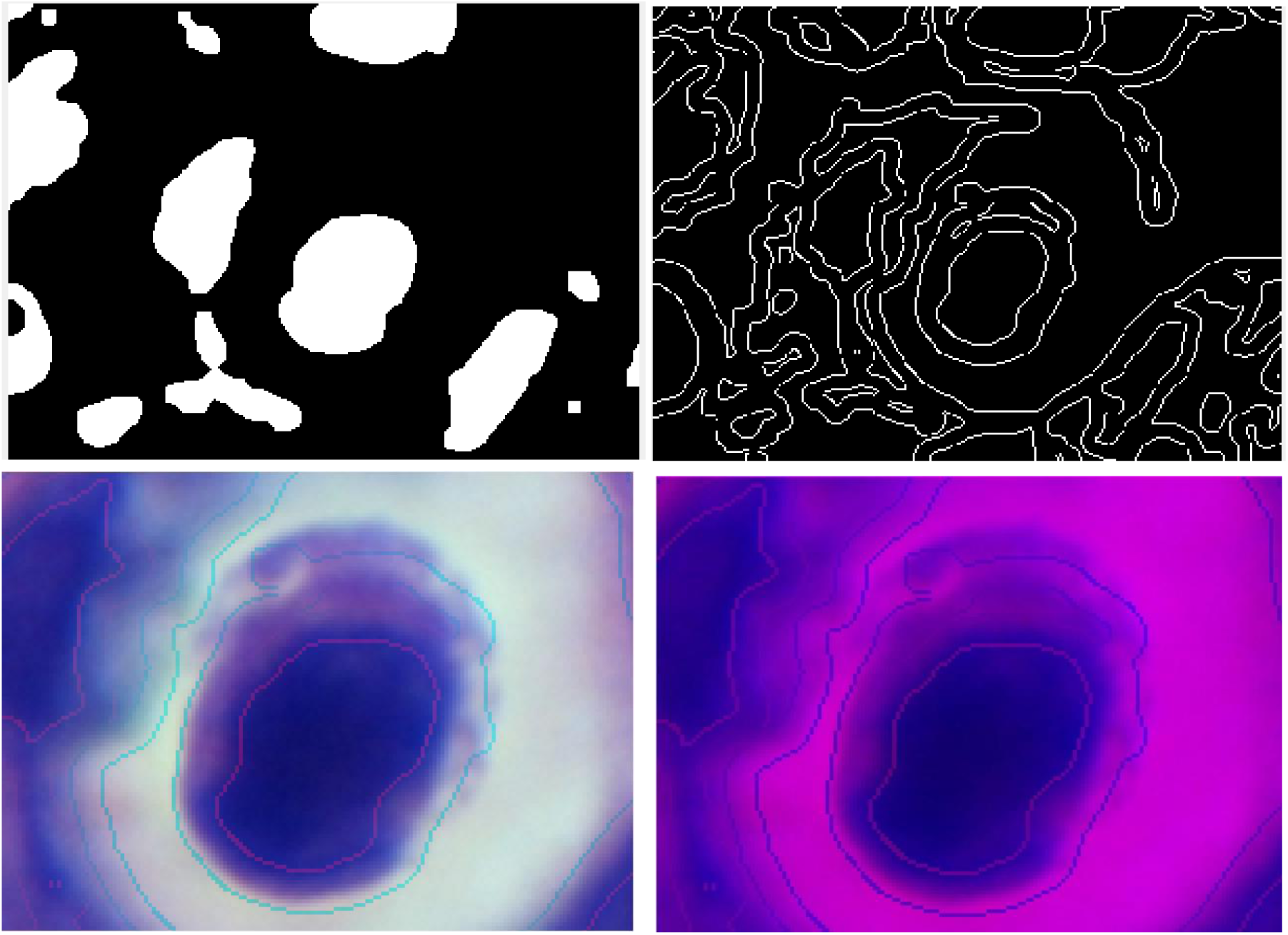

**Figure 4.**
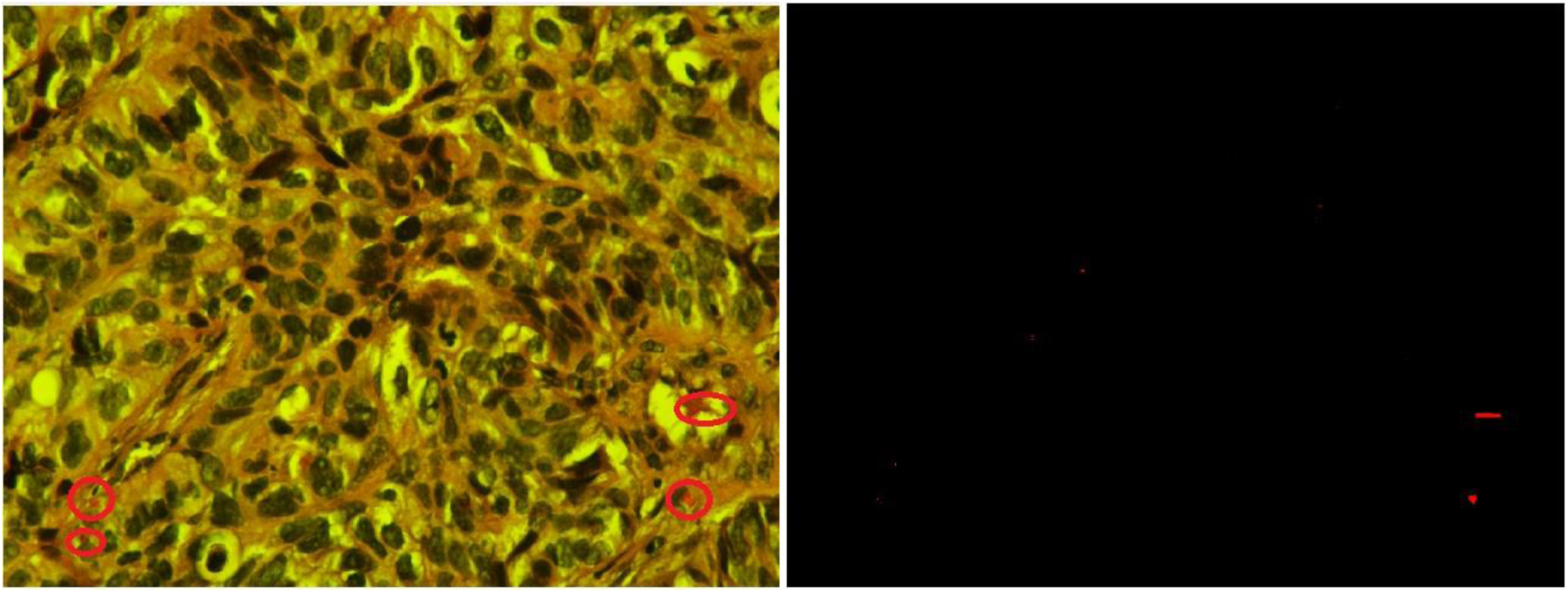

### E. Alignment an cell polarization pattern detection

Using the detected nuclei and cytoplasm in step A, it is possible to measure the orientation based on the angle of rotation of an ellipse formula around it. Using the vector rotation of Quadrants II and IV as positive and I and III as negative, obtaining an overall summation and calculating and index of orientation based on each individual cell. (fig 5)

**Figure 5.**
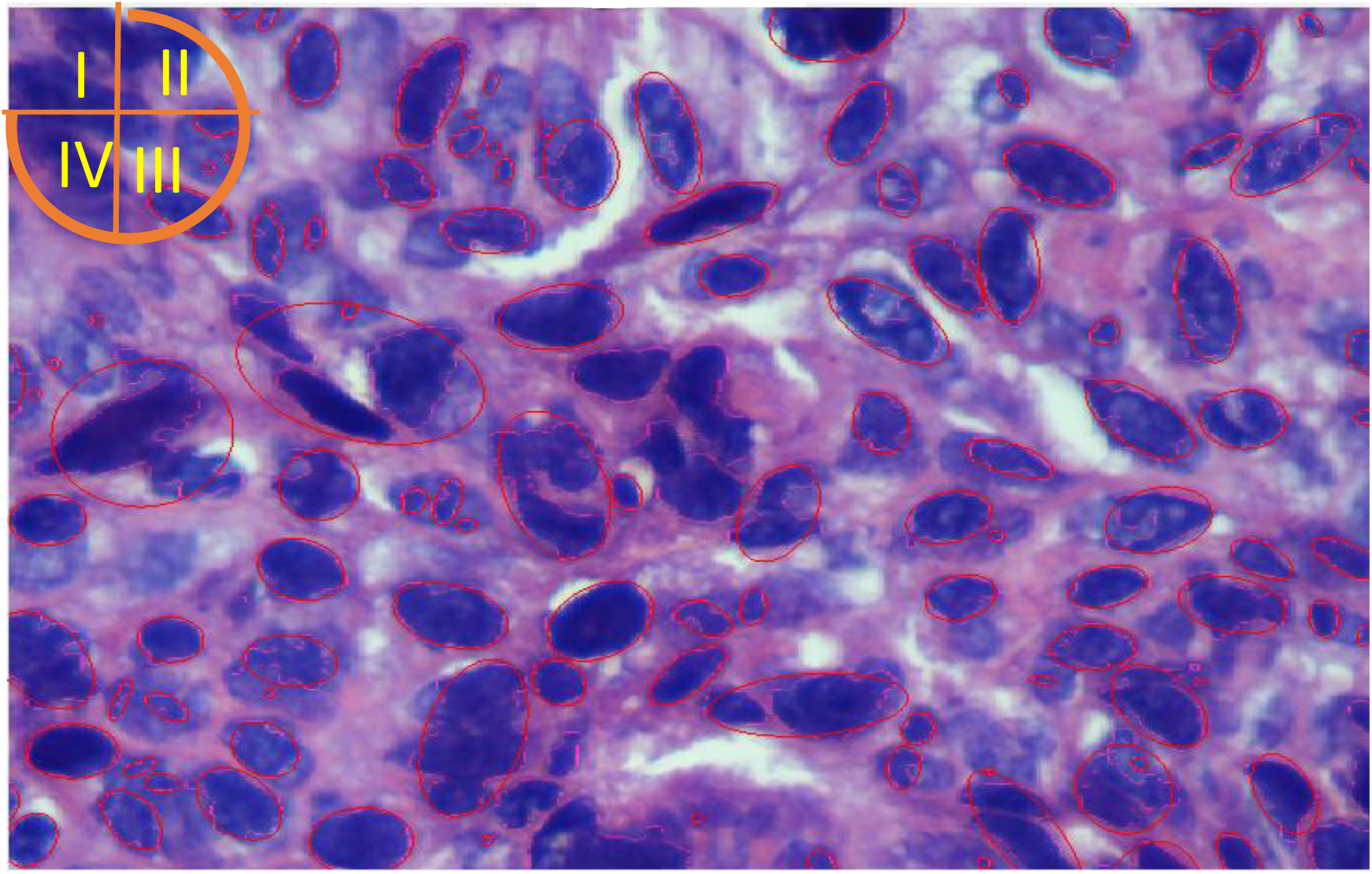

### F. Increment of Mitotic cell activity detection

Using the Ellipse function, it is calculated the area inside the delimited section, and calculated the void volume near to the center of the ellipse. This method only works around 30% of times, with mitotic figures in dividing process contrary to a more robust method like proposed by Saha et al. (4). (Figure 6)

**Figure 6.**
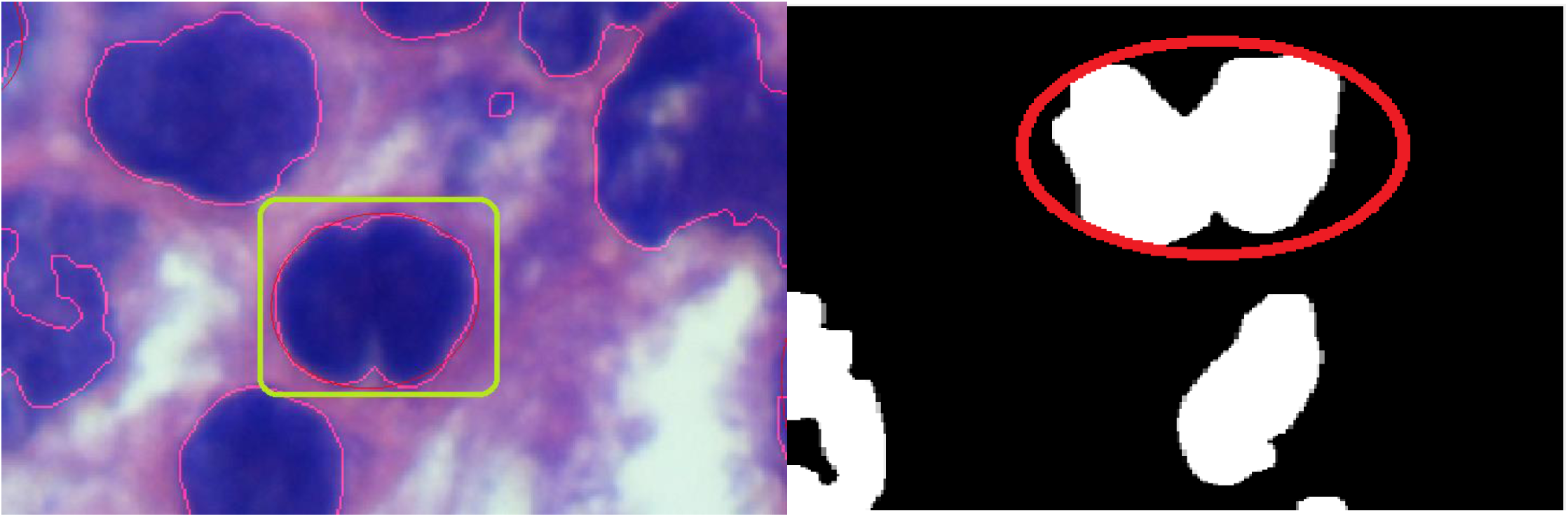

**Figure 7:**
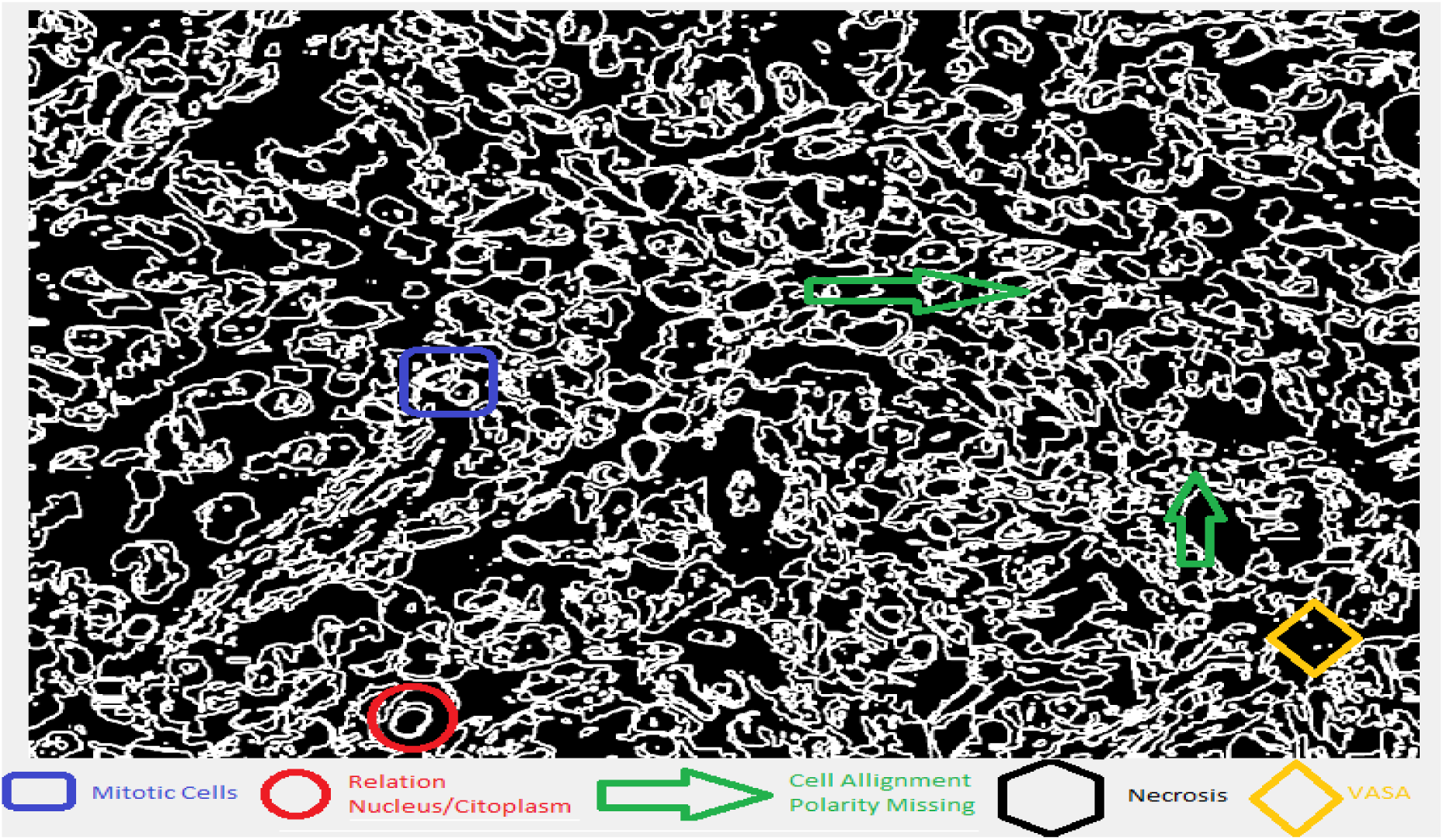
Expected Results.

**Figure 8:**
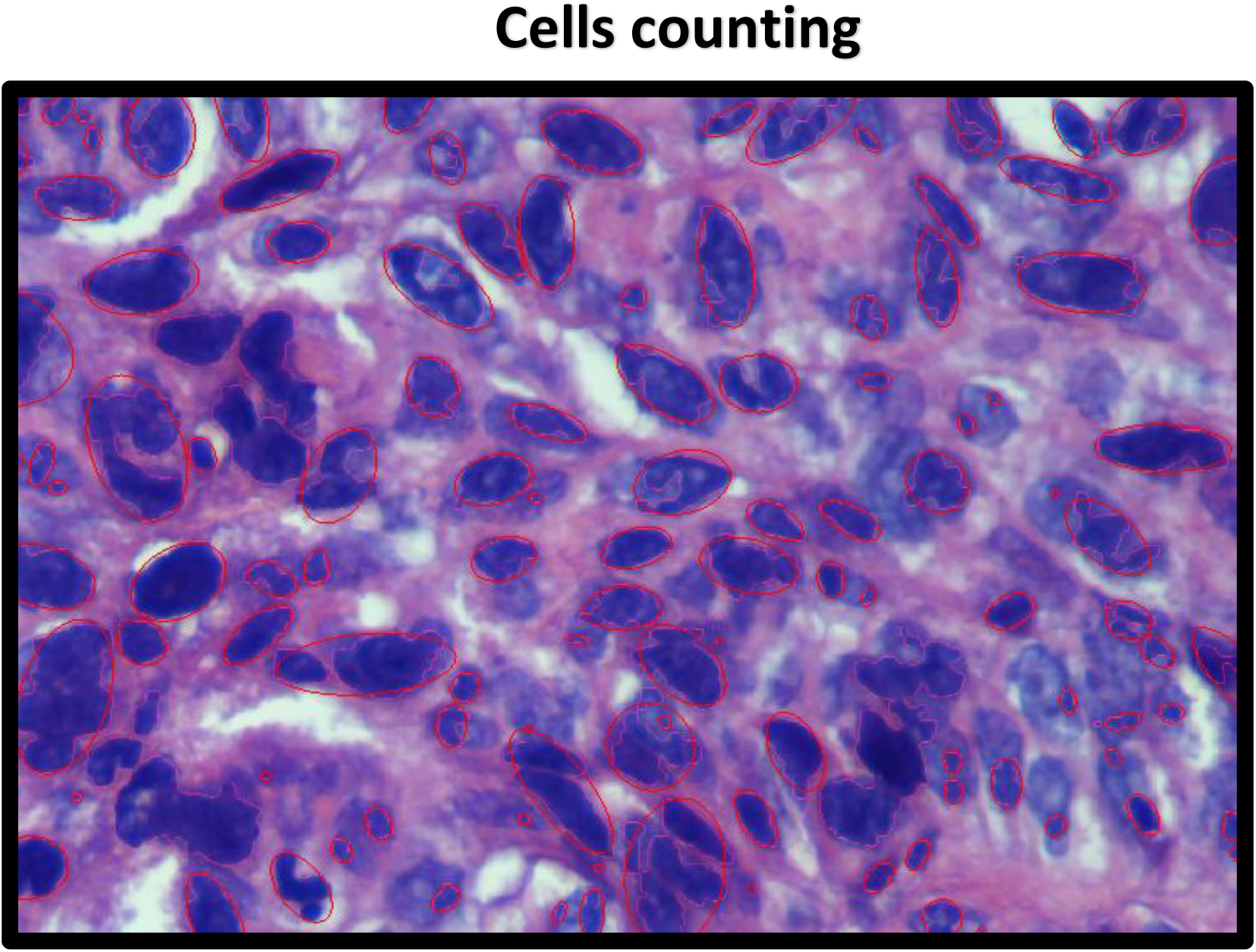
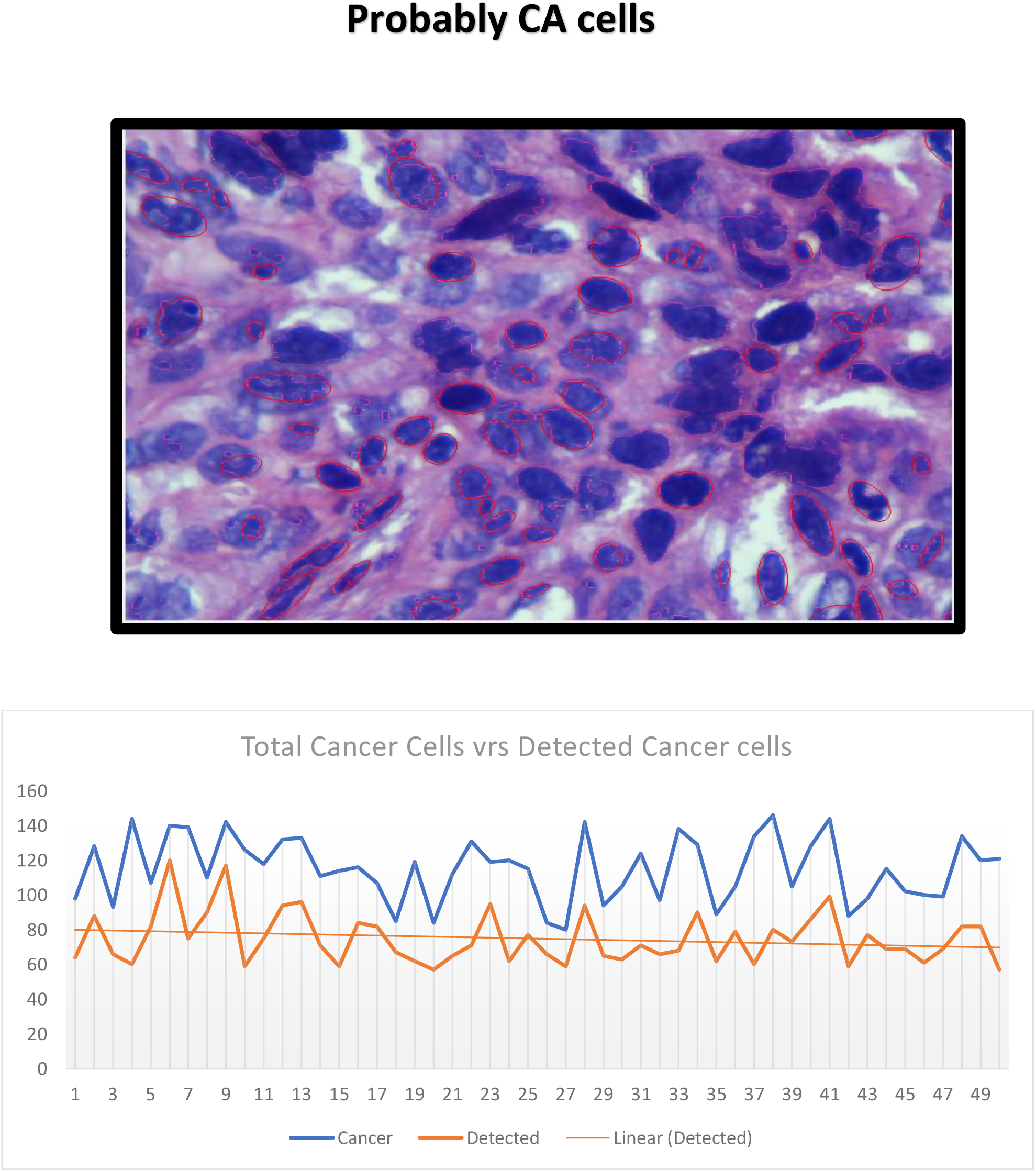
Obtained Results Stage II tissue.

| Sample | Cancer | Detected | Percentage | Sample | Cancer | Detected | Percentage |
| --- | --- | --- | --- | --- | --- | --- | --- |
| 1 | 98 | 64 | 65% | 26 | 84 | 66 | 79% |
| 2 | 128 | 88 | 69% | 27 | 80 | 59 | 74% |
| 3 | 93 | 66 | 71% | 28 | 142 | 94 | 66% |
| 4 | 144 | 60 | 42% | 29 | 94 | 65 | 69% |
| 5 | 107 | 82 | 77% | 30 | 105 | 63 | 60% |
| 6 | 140 | 120 | 86% | 31 | 124 | 71 | 57% |
| 7 | 139 | 75 | 54% | 32 | 97 | 66 | 68% |
| 8 | 110 | 90 | 82% | 33 | 138 | 68 | 49% |
| 9 | 142 | 117 | 82% | 34 | 129 | 90 | 70% |
| 10 | 126 | 59 | 47% | 35 | 89 | 62 | 70% |
| 11 | 118 | 75 | 64% | 36 | 105 | 79 | 75% |
| 12 | 132 | 94 | 71% | 37 | 134 | 60 | 45% |
| 13 | 133 | 96 | 72% | 38 | 146 | 80 | 55% |
| 14 | 111 | 71 | 64% | 39 | 105 | 73 | 70% |
| 15 | 114 | 59 | 52% | 40 | 128 | 86 | 67% |
| 16 | 116 | 84 | 72% | 41 | 144 | 99 | 69% |
| 17 | 107 | 82 | 77% | 42 | 88 | 59 | 67% |
| 18 | 85 | 67 | 79% | 43 | 98 | 77 | 79% |
| 19 | 119 | 62 | 52% | 44 | 115 | 69 | 60% |
| 20 | 84 | 57 | 68% | 45 | 102 | 69 | 68% |
| 21 | 112 | 65 | 58% | 46 | 100 | 61 | 61% |
| 22 | 131 | 71 | 54% | 47 | 99 | 69 | 70% |
| 23 | 119 | 95 | 80% | 48 | 134 | 82 | 61% |
| 24 | 120 | 62 | 52% | 49 | 120 | 82 | 68% |
| 25 | 115 | 77 | 67% | 50 | 121 | 57 | 47% |
|  |  |  |  | Average | 117.72 | 77.52 | 66% |

## III. Results

After perform step A to F in 50 different images of poorly differentiated adenocarcinoma in stages 3 and 4. It is possible to assure that the method for detect nuclei and cytoplasm in cells in this specific images varies from 85% to 95%, taking on count that if cells volume or cells pixels is inferior to 200, could not be detected.

This measure provides and index involving the relationship between this two components, being consider as normal when this relationship is where the cytoplasm is two or 3 times the nuclei, if this relationship is inverted this cell was added to the pool as possible cancer cells, finally the total count of cells was divided between the cell considered as normal against the cancer cells. The average index in the 50 samples was 0.14, being 1 as total normal tissue, and 0 a tissue completely formed by cancer cells.

But because the lack of reliability on just one parameter for measuring something as important as cancer in tissue, is necessary to add more parameters in this detection process.

The next parameter in order of importance was the alignment and cell orientation pattern, as mentioned in step E, the image was divide in four Quadrants and the polarity was measured as +1 (positive, Quadrants II, IV) and -1 (negative, Quadrants I, III), the orientation was then measured as more than 70% aligned in the same direction as completely normal, and ranges from 40% to 69% as misaligned pattern (possible cancer pattern).

The third parameter in order of importance was the mitotic figures or the abnormal number and presence of those in a cellular tissue, the estimated presence in non-cancer sample is around 20%, so the program was designed to find the volume inside the elliptical area, as described in step F, but even after several trials, the program was only able to detect around 15% to 25% of the mitotic cells, not being enough reliable to take in consideration as cancer detection parameter.

The two last parameters measured were the Vasculature and necrosis presence on tissue, from the 50 images, only 3 got presence of necrosis and was not enough to perform enough testing on the analysis but from this 3 images, the necrotic area detected was around 90%. In accordance to the vascular analysis and the increment of that, as shown by figure 4, the program detect around 95% in the fifty images. But the presence of vascularity in this tissue based on the type of cancer selected, lack of significance.

As final conclusion, using this program will give you a percentage of 60% to 70% of chances on select tissue with cancer cells, for further analysis by a trained pathologist. This coding software could work as a screening when the number of images to analyze or time for observing those imager, makes difficult the work for one observer.

It is important to remark, that a posterior analysis comparing normal tissue against cancer cells and different stages will be necessary to probe or validate its clinical use, in this moment that comparison is not possible based on the time for this developing, and also that the department of Pathology of Guatemala don’t possess a database on normal tissue, only pathological ones. Making necessary to create a protocol on a prospective analysis.

In the samples table, it is assumed that every counted cells were a cancer cell, because it was sampled from stage 4 cancer tissue, however to perform and accurate measure it will be necessary to do a statistical analysis, where every cell marked as a detected will be process to determinate if it was really a cancer cell.

This further analysis will be then able to compare the prognosis against the real diagnostic in cell by cell population cloning.

